# Uncovering bacterial hosts of class 1 integrons in an urban coastal aquatic environment with a single-cell fusion-polymerase chain reaction technology

**DOI:** 10.1101/2023.02.09.527782

**Authors:** Qin Qi, Timothy M Ghaly, Anahit Penesyan, Vaheesan Rajabal, Jeremy AC Stacey, Sasha G Tetu, Michael R Gillings

## Abstract

Horizontal gene transfer (HGT) is a key driver of bacterial evolution via transmission of genetic materials across taxa. Class 1 integrons are genetic elements that correlate strongly with anthropogenic pollution and contribute to the spread of antimicrobial resistance (AMR) genes via HGT. Despite their significance to human health, there is a shortage of robust, culture-free surveillance technologies for identifying uncultivated environmental taxa that harbour class 1 integrons. We developed a modified version of epicPCR (<u>e</u>mulsion, <u>p</u>aired <u>i</u>solation and <u>c</u>oncatenation <u>p</u>olymerase <u>c</u>hain <u>r</u>eaction) that links class 1 integrons amplified from single bacterial cells to taxonomic markers from the same cells in emulsified aqueous droplets. Using this single-cell genomic approach and Nanopore sequencing, we successfully assigned class 1 integron gene cassette arrays containing mostly AMR genes to their hosts in coastal water samples that were affected by pollution. Our work presents the first application of epicPCR for targeting variable, multi-gene loci of interest. We also identified the *Rhizobacter* genus as novel hosts of class 1 integrons. These findings establish epicPCR as a powerful tool for linking taxa to class 1 integrons in environmental bacterial communities and offer the potential to direct mitigation efforts towards hotspots of class 1 integron-mediated dissemination of AMR.

**Synopsis:** We present a novel single-cell genomic surveillance technology for identifying environmental bacterial hosts of a class of mobile genetic elements that are linked to anthropogenic pollution and contribute to the dissemination of antimicrobial resistance.

## Introduction

Horizontal gene transfer (HGT) is an important source of genomic variability upon which selection can act ^1–3^. Microbial adaptation via HGT most commonly occurs among closely related taxa or between more distantly related taxa found in similar environments ^4, 5^. A key example is the dissemination of antimicrobial resistance (AMR) genes via conjugation, bacteriophage transduction, and natural transformation by extracellular DNA ^6–9^. To understand the dynamics of HGT, we need to identify the sources of horizontally transferred genes and the range of taxa they subsequently inhabit ^10–12^. This is challenging given the sheer diversity of microbial taxa and the difficulty of culturing many environmental microbes ^13–16^. Metagenomic sequencing of bacterial communities does not require prior culturing of microbes ^17, 18^. However, mobile genetic elements are often associated with repetitive DNA sequences and exhibit complex interactions with genome rearrangements, which limit our ability to attribute laterally transferred genes to host genomes due to assembly and/or taxonomic binning difficulties ^19, 20^.

epicPCR (<u>e</u>mulsion, <u>p</u>aired <u>i</u>solation and <u>c</u>oncatenation PCR) is a fusion PCR-based technique that overcomes these problems by co-amplifying taxonomic markers with functional genes of interest from single uncultured bacterial cells. Single cells are trapped in droplets generated by emulsifying an aqueous phase containing PCR reagents and environmental bacteria in an emulsion oil ^21–26^. This compartmentalises the fusion PCR into parallel, single cell-level reactions. Within each droplet, chimeric amplicons that combine genes of interest with taxonomic markers corresponding to diverse phylogenetic groups are generated for high-throughput sequencing. To date, epicPCR has been successfully applied to target functional genes including dissimilatory sulfite reductase *dsrB* genes ^21, 24^, antibiotic resistance genes ^22, 23, 26, 27^, class 1 integron associated genes (*intI1, sul1*, Δ*qacE*) ^23, 27, 28^, and bacteriophage-borne ribonucleotide reductase *RNR* genes ^25^. epicPCR amplicons typically comprise short fragments of genes of interest linked to the V4 hypervariable region of 16S rRNA markers. Illumina sequencing of recovered amplicons demonstrates the presence of the genes of interest in a variety of bacterial taxa. However, no studies have so far applied epicPCR to target variable, multi-gene loci of interest.

Class 1 integrons contribute to the dissemination of AMR genes via HGT in natural ecosystems and clinical settings ^29, 30^. Integrons are genetic elements that comprise an integron-integrase gene (*intI*), an *attI* recombination site, and a promoter (P_C_) that drives the expression of variable gene cassettes ^31–33^. IntI catalyses the integration and excision of gene cassettes via *attC* sites that can be recombined with *attI* or other *attC* sites ^33–36^. All class 1 integrons contain highly conserved integron-integrase (*intI1*) and *attI1* sequences (Figure 1A). Clinical class 1 integrons are strongly associated with AMR gene cassettes that confer resistance to antibiotics, disinfectants, or heavy metals ^29, 31, 37^. Mobile genetic elements such as broad host-range plasmids and transposons have acquired class 1 integrons, resulting in the proliferation of class 1 integrons across diverse bacterial hosts ^31, 38^. Recently, it has been reported that phage-plasmids can also disseminate class 1 integrons ^39^.

**Figure 1.**
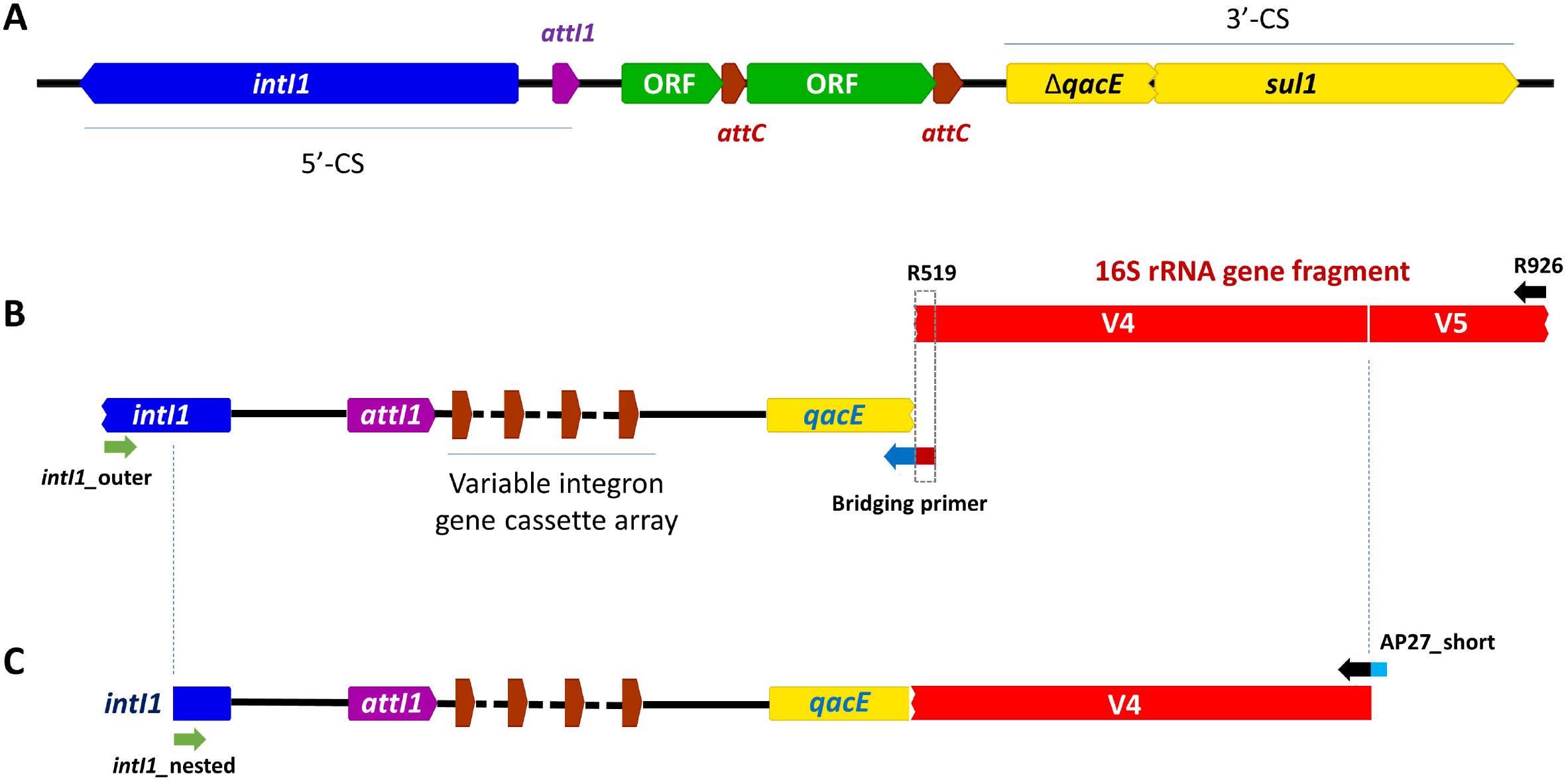
Class 1 integrons and their co-amplification with 16S rRNA marker genes via epicPCR. **(A)** The class 1 integron carried by the IncW plasmid R388 has the canonical structure of clinical class 1 integrons. *intI1* and *attI1* sequences are highly conserved in the 5’-conserved segments (CS). The 3’-CS comprise a truncated quaternary ammonium compound (QAC) resistance gene (Δ*qacE*) and a sulfonamide resistance gene (*sul1*). Between the 5’-CS and 3’-CS are variable integron gene cassette arrays that contain *attC* recombination sites and gene cassettes. Each gene cassette is typically associated with a single open-reading frame (ORF). **(B)** During the first stage of epicPCR, the *intI1*_outer primer and the R519_*qacE* bridging primer target *intI1* and *qacE* sequences in the 5’-CS and 3’-CS respectively and amplify the class 1 integron. A short overhang introduced by the bridging primer to the *qacE* fragment end allows the intermediate PCR product to act as a long primer and co-amplify the V4-V5 hypervariable regions of the 16S rRNA marker gene in the presence of the R926 reverse primer. **(C)** During the second stage of epicPCR, the *intI1*_nested and AP27_short nested primers further amplify the chimeric amplicons pooled from the previous step. AP27_short introduces a non-degenerate tail sequence (in blue) to the 3’-end of the amplicons. During standard PCR verification of chimeric amplicon formation (Figure S3A), this tail sequence is targeted by the AP28_short non-degenerate primer, which is used in conjunction with the *qacE*_F forward primer. The final chimeric epicPCR amplicons contain highly conserved features of class 1 integrons, including a 157 bp *intI1-attI1* region in the 5’-CS and the *qacE*/Δ*qacE* fragment in the 3’-CS.

Bacterial hosts of class 1 integrons have previously been isolated and characterised through colony PCR screening and fosmid library preparation ^29, 40–43^. The necessity to pre-isolate environmental bacteria severely biases host identification towards the cultivated fraction and hinders our ability to attribute relative abundances of class 1 integrons to individual taxa. To overcome these limitations, we developed a modified experimental and bioinformatic pipeline for epicPCR that associates class 1 integrons with their bacterial hosts. Class 1 integron gene cassette arrays are co-amplified with the V4 hypervariable region of the 16S rRNA marker genes from individual bacterial cells (Figure 1B-C). The fused amplicons are sequenced using Nanopore long-read technology. We applied stringent bioinformatic filtering to ensure that the associations between the class 1 integrons and the taxonomic markers were *bona fide*. We successfully tested our epicPCR pipeline on water samples from a coastal environment and linked six unique class 1 integrons with variable gene cassettes to seven distinct environmental bacterial taxa. While most of the gene cassettes encode AMR genes, we also linked a gene cassette encoding a cupin-domain containing protein of unknown functions to two *Alphaproteobacteria* hosts. Furthermore, our results demonstrated that *Rhizobacter* is a novel host of class 1 integrons. Overall, our study shows that epicPCR is a powerful tool for determining the environmental hosts of class 1 integrons with the potential to access previously untapped carriers of these genetic elements. This paves the way for future studies that seek to pinpoint the “sources” and “sinks” of class 1 integrons.

## Materials and Methods

### Isolation of bacteria from an urban coastal location

Coastal water samples were collected in three 50 mL biological replicates after rain in a rocky intertidal zone downstream from a stormwater outlet at Shark Point (location coordinates: - 33.91, 151.27) near Sydney, NSW, Australia (Figure S1). The samples were filtered using EASYstrainer cell strainers with a mesh size of 40 μm (Greiner AG, Germany) and centrifuged at 3220 *g* for 15 minutes. Cell pellets were resuspended in phosphate buffered saline (PBS) solution at pH 7.4. Resuspended cells were stored as 25% glycerol stocks.

### Bacterial cell count estimates by fluorescence microscopy

Approximately 10^5^-10^6^ cells were fixed and stained in PBS containing 4% formaldehyde and 2 μg/mL 4’,6-diamidino-2-phenylindole (DAPI) for 30 min in the dark with rotation at room temperature. Re-suspended cells were diluted in PBS, transferred to a haemocytometer and imaged under the x40 lens of an Olympus BX63 fluorescence microscope. Differential interference contrast (DIC) and DAPI fluorescence channel images were overlaid and analysed with ImageJ (National Institutes of Health, United States). Bacterial cell densities were calculated from the average number of cells in five 0.2 mm x 0.2 mm square grids of the counting chamber for each sample (Figure S2).

### Emulsion, paired-isolation and concatenation PCR (epicPCR)

Our hybrid epicPCR procedures were adapted from the protocols of Sakowski *et al* ^25^ and Diebold *et al* ^26^. Briefly, epicPCR was performed in three technical replicates for three biological replicates. Approximately 150,000 cells from the bacterial glycerol stocks were pelleted and vortexed in a 75 μL PCR reagent mix containing GC buffer (1x), dNTP mix (0.4 mM), Phusion Hot-Start II polymerase (0.05 U/μL) (Thermo Scientific, United States), bovine serum albumin solution (1 μg/μL) (Promega, United States), Lucigen Ready-Lyse lysozyme (500 U/μL) (LGC Biosearch Technologies, United States) and oligonucleotide primers R926 (2 μM), *intI1*_outer (1 μM), and R519-*qacE* bridging primer (0.04 μM) (Table S2). To demonstrate the absence of false associations between 16S rRNA markers and class 1 integrons originating from different cells, we spiked one set of technical replicates with cells of a class 1 integron-free *Escherichia coli* MG1655 strain at a total population frequency of approximately 10%.

ABIL emulsion oil was prepared by supplementing mineral oil with 4% ABIL EM 90 (Redox, Australia) and 0.05% Triton X-100 (Promega, United States). 425 μL of ABIL oil was added to the PCR mix, which was immediately emulsified at 4 ms^−1^ for 45 s using the FastPrep-24 bead beating system (MP Biomedicals, United States). The water-in-oil emulsion was aliquoted into 8 portions, incubated at 37°C for 10 min, and was subjected to the following PCR amplification conditions: 98°C for 5 minutes; 38 cycles of 98°C for 10 s, 59°C for 20 s, and 72°C for 2 min; and finally 72°C for 5 min. The *intI1*_outer primer and the R519-*qacE* bridging primer bind to *intI1* in the 5’-conserved segment (CS) and *qacE* in the 3’-CS of class 1 integrons respectively (Figure 1B). The 18 bp overhang introduced by the bridging primer to the *qacE* end allows the intermediate PCR products to act as a long primer and amplify the V4-V5 hypervariable region with the R926 reverse primer.

The emulsified PCR mix for each technical replicate was pooled, vortexed in 900 μL of isobutanol and 200 μL of 5M NaCl and centrifuged for 1 min at ~20,000*g* with a soft brake. The aqueous phase at the bottom of the tubes was extracted, purified using the Monarch PCR & DNA Cleanup Kit (New England Biolabs, United States), and eluted with nuclease-free water. Sera-Mag Select magnetic beads (Cytiva, United States) were added to the eluted PCR products to deplete <300 bp DNA fragments. The size-selected DNA was used as template in 100 μL Phusion PCR mix containing GC buffer (1x), dNTP mix (0.4 mM), Phusion Hot-Start II polymerase (0.04 U/μL), AP27_short primer (0.8 μM), *intI1*_nested primer (0.4 μM), forward and reverse blocking primers (0.32 μM each). The PCR mix was subjected to the following thermocycling conditions in 8 aliquots: 98°C for 30 s; 35 cycles of 98°C for 10 s, 59°C for 20 s and 72°C for 1 min 50 s; followed by a final step of 72°C for 5 min. epicPCR products were visualised on 1% agarose gel by electrophoresis and treated with Sera-Mag Select magnetic beads to deplete epicPCR products containing cassette-less integrons.

To confirm that fusion between class 1 integron and 16S DNA fragments has occurred (Figure S3A), 2 ng of each purified epicPCR product was amplified using *qacE*_F (0.4 μM) and AP28_short (0.4 μM) primers in GoTaq polymerase master mix (Promega, United States) under the following thermocycling conditions: 95°C for 30 s; 30 cycles of 95°C for 30 s, 55°C for 30 s and 72°C for 30 s; with a final step of 72°C for 5 min. Successful amplification of the *qacE-16S* rRNA gene chimeric region should produce ~390 bp gel bands.

### Sanger sequencing of epicPCR products from the *Rhizobacter* genus

To confirm that class 1 integrons can be found in the *Rhizobacter* genus, we pooled approximately 150,000 cells from the three biological replicates and performed the epicPCR procedures in three technical replicates with the minor modifications shown in Figure S3B: the R926 and AP27_short primers were replaced with *Rhizobacter*-specific primers *Rhizobacter_16S*_ R926 and *Rhizobacter_16S*_ R806 respectively, which were designed based on the 16S rRNA sequence of *R. gummiphilus* strain NS21 (NCBI accession no. CP015118) ^44^. *qacE*_F was used as the nested forward primer. The resulting 389 bp epicPCR products were verified by Sanger sequencing (Marcogen Inc, South Korea).

### Oxford Nanopore long-read sequencing

Purified epicPCR amplicons were treated with the NEBNext Ultra II End Repair/dA-Tailing Module (New England Biolabs, United States), purified with JetSeq Clean magnetic beads (Meridian Bioscience, United States), and eluted using nuclease-free water. End-repaired DNA was barcoded using the Native Barcoding Expansion kit (ONT, United Kingdom) and Blunt/TA Ligase Master Mix (New England Biolabs, United States). Barcoded samples were multiplexed and ligated to sequencing adaptor molecules in Adapter Mix II using the NEBNext Quick T4 DNA ligase (New England Biolabs, United States). The ligation products were washed twice with Short Fragmentation Buffer and eluted with Elution Buffer. The sequencing library was loaded into a FLO-MIN106D flow cell R9.4.1 and a MinION Mk1B sequencer (ONT, United Kingdom). The minimum threshold for read length was set to 1 kB on the MinKNOW operating software. Basecalling of FAST5 raw data was performed with the high accuracy option using Guppy v6.1.2 (ONT, United Kingdom). The native barcoded samples were demultiplexed using default parameters.

### Analysis of Nanopore sequencing data of epicPCR amplicons

All filtering and processing steps were carried out using an in-house pipeline (https://github.com/timghaly/Int1-epicPCR). First, the pipeline quality-filters reads using NanoFilt v2.8.0 ^45^, removing those with an average read quality of <7 and read length <670 bp, which represents the minimum length of an epicPCR product with a cassette-less integron. Quality-filtered reads are then oriented and trimmed with the final nested forward and reverse primer sequences using Pychopper v2.7.0 (https://github.com/epi2me-labs/pychopper). Pychopper identifies both primers in each read using edlib v1.2.3 ^46^, orients the reads based on the forward and reverse primer sequences, and discards reads that do not contain both primers in the correct orientations. The pipeline then clusters the primer-oriented reads into amplicon-specific clusters using isONclust v0.0.6.1 ^47^. Error correction was performed on each cluster using isONcorrect v0.0.8 ^48^, which can jointly use all cassette arrangements of the same integron that occur in different clusters and allow efficient error correction even for amplicons with low sequencing depths. A consensus sequence is then generated for each cluster using spoa v4.0.7 ^49^. All consensus sequences are then pooled, while removing any redundancies, including reverse complement redundancies, using dedupe from BBTools v35 (https://github.com/kbaseapps/BBTools). Next, the consensus sequences are screened for the R519-*qacE* bridging primer and the 157 bp region (covering the 5’-end of *intI1* and all of *attI1*) using blastn from BLAST v2.2.31. Retained sequences must contain both regions. Any sequences containing more than one hit to either of these sequences are considered unintended chimeras and discarded. Finally, sequences are screened for the correctly fused 16S rRNA gene fragment using Metaxa2 v2.2.3 ^50^. The final output of the pipeline for each sample is a set of full-length, primer-oriented amplicon consensus sequences that contain a complete *attI1* sequence, R519-*qacE* bridging primer, and the V4 hypervariable region of the 16S rRNA gene. Cd-hit v4.8.1 ^51^ was used to cluster chimeric epicPCR products comprising class 1 integron gene cassette arrays and V4 hypervariable regions of the 16S rRNA gene with ≥99% pairwise identity in nucleotide sequences in at least 3 epicPCR replicates. For epicPCR products that were found in fewer than 3 replicates, the NCBI Genome Workbench was used to perform blastn searches of these sequences against a local database created using all the Nanopore reads obtained in this study. Reads with ≥98% pairwise identity in nucleotide sequences and ≥98% coverage were aligned using the Geneious bioinformatic software (Biomatters, New Zealand), and the consensus sequences were generated from these alignments within each set of epicPCR replicates. Consensus sequences that could be found in at least 3 epicPCR replicates and showed ≥99% pairwise identity in nucleotide sequences and ≥99% identical sites were added to the list of epicPCR products that had already been shortlisted by the Cd-hit algorithm. IntegronFinder 2.0 [parameters: --local-max --gbk --promoter-attI --calin-threshold 1] was used to predict the ORFs and *attC* sites ^52^.

### Taxonomic classification for bacterial hosts of class 1 integrons

The sequences of V4 hypervariable regions of 16S rRNA gene sequences obtained in this study were searched against the SILVA 16S rRNA gene database website ^53^ using the Alignment, Classification and Tree (ACT) tool with a cut-off threshold of 0.80. The V4 region sequences were aligned using the Geneious multiple alignment tool using default parameters and cost matrix = 93% similarity. A phylogenetic tree was constructed with PhyML (v3.0) using the maximum likelihood method based on the Hasegawa-Kishino-Yano (HKY85) model with bootstrapping (number of replicates = 100) ^54^. To verify genus-level taxonomic classifications, the V4 hypervariable regions of 16S rRNA for the *Rhizobacter* and *Aquabacterium*-like hosts found in this study were searched against the NCBI Nucleotide Collection by blastn with the exclusion of “uncultured/environmental sample sequences”. The 16S rRNA sequences of all identical matches were searched against the SILVA 16S rRNA gene database. All the reference genomes of *Rhizobacter* and *Aquabacterium* species were downloaded from the NCBI RefSeq database (last accessed on 11 November 2022). Nucleotide alignment was performed against the full 16S rRNA gene sequences from the RefSeq genomes using the Geneious bioinformatic software.

## Results and Discussion

Coastal water samples were collected in three biological replicates from the rockpools downstream of a stormwater outlet at Shark Point, Sydney, New South Wales, Australia (Figure S1). We anticipated that class 1 integron-associated environmental bacteria would be detected at this location, because surface runoff was expected to carry pollutants from the surrounding urban areas. A preponderance of cells displayed unicellular and planktonic morphologies (Figure S2). We applied modified epicPCR procedures to amplify class 1 integron gene cassette arrays from single bacterial cells encapsulated by PCR reagent droplets homogenously dispersed in an oil emulsion and successfully generated chimeric epicPCR products containing class 1 integrons and 16S rRNA gene sequences (Figure 1). These epicPCR products were Nanopore sequenced. The median number of reads per sample was 232,402. After stringent bioinformatic filtering, we identified six class 1 integrons in seven distinct bacterial hosts (Figure 2). All seven combinations of epicPCR products were observed in at least three independent replicates (*n*≥3, ≥99% pairwise nucleotide identity and ≥99% identical sites relative to the consensus sequence for each epicPCR product). In addition, there were five combinations of class 1 integrons and hosts that were detected in fewer than 3 replicates and were consequently excluded. Taxonomic classifications were obtained using the SILVA ribosomal RNA gene database (Table S1) ^53^. *Alphaproteobacteria* (*n*=2) and *Gammaproteobacteria* (*n*=5) were the two major classes of hosts represented in our dataset, while *Burkholderiales* (*n*=4) was the most common bacterial order. These observations are broadly consistent with previous findings that class 1 integrons are associated with *Gammaproteobacteria* ^29, 42, 43^. The finding of integrons in two *Alphaproteobacteria* is notable, because *Alphaproteobacteria* are rarely known to be associated with integrons ^55^. Currently, there are very few examples of *Alphaproteobacteria* species that harbour class 1 integrons ^56–58^. This underscores the potential of epicPCR approaches to discover novel host-function associations.

**Figure 2.**
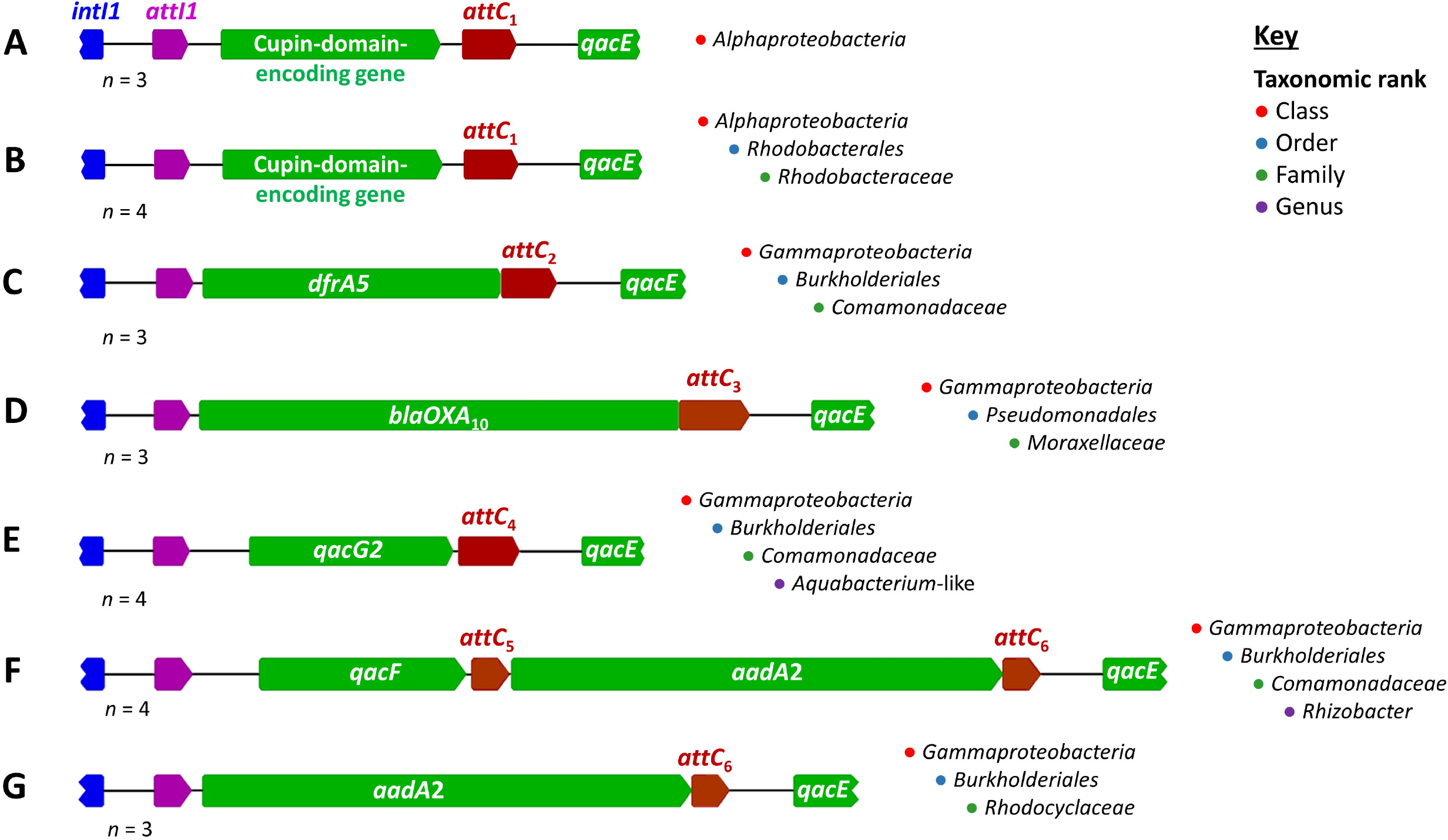
Class 1 integrons identified through epicPCR alongside the taxonomic classification of their respective bacterial hosts. The *intI1* gene fragment is displayed in blue, the *attI1* site in purple, and all the gene cassette ORFs in green. The six different *attC* sites are shown in red.

Six different gene cassettes were detected, five of which encoded well-characterised AMR genes (Table 1). The nucleotide sequences of all the gene cassettes reported in this study are summarised in Table S3. We compared their nucleotide sequences against reference class 1 integron gene cassettes from the NCBI database and found that both the coding and non-coding regions of these gene cassettes were highly conserved relative to the reference sequences ^34, 59^. In our dataset, there are six different *attC* recombination sites with 100% nucleotide sequence identity to those in the respective reference gene cassettes in Table 1. We also aligned all the nucleotide sequences in the 5’-CS and 3’-CS regions of all the class 1 integrons from this study and found that both the 157 bp *intI1/attI1* and 190 bp *attC/qacE* regions in the chimeric amplicons were identical in all but one exception (Figure S4). Overall, these class 1 integron structures are typical of those recovered from clinical bacterial isolates ^29, 60, 61^.

**Table 1.** Comparison between the class 1 integron gene cassettes sequenced in this work and annotated reference class 1 integron gene cassettes.

| ORF in gene cassette | Predicted function of ORF gene product | NCBI reference gene cassette | % Nucleotide identity to NCBI reference gene cassette sequence |
| --- | --- | --- | --- |
| Cupin-domain-encoding gene | Unknown | FJ172388 | 99.8% nucleotide identity; 1 synonymous mutation in the coding region |
| *dfrA5* | Trimethoprim-resistant dihydrofolate reductase | X12868 | 100% nucleotide identity |
| *blaOXA10* | Oxacillin-hydrolysing beta-lactamase | XXU37105 |  |
| *qacG2* | Quaternary ammonium compound efflux pump | AF327731 | 99.6% nucleotide identity; 2 non-synonymous mutations (S13F and A57G) in the coding region |
| *qacF* | Quaternary ammonium compound efflux pump | AF034958 | 99.6% nucleotide identity; 1 non-synonymous mutation (S10A) in the coding region |
| *aadA2* | Aminoglycoside (3'') adenylyltransferase conferring resistance to streptomycin and spectinomycin | X68227 | In the *Rhizobacter* host: 100% nucleotide identity |
|  |  |  | In the *Rhodocyclaceae* host: 99.6% nucleotide identity; 1 SNP in the non-coding region, 1 non-synonymous mutation (V257A) in the coding region |

### Class 1 gene cassettes detected by epicPCR

Cupins belong to a superfamily of proteins with a beta-barrel fold and were originally found as highly conserved domains in plant proteins ^62^. We found a gene cassette encoding a cupin barrel domain of unknown functions in two *Alphaproteobacteria* hosts. One host was unclassified at the order level (Figure 2A), while the other belonged to the *Rhodobacteraceae* family (Figure 2B). This cupin-domain-encoding gene was previously observed in a class 1 integron gene cassette from an unknown host ^59^. Based on identical nucleotide sequence matches from the NCBI nucleotide database, this gene cassette was detected in class 1 integrons, non-class 1 integrons, and CALINs (clusters of *attC*s lacking an associated integron-integrase) in the chromosomes or plasmids of several *Gammaproteobacteria* hosts (Table S4). This observation suggests selective pressure for maintaining this gene cassette in both class 1 and non-class 1 integron genetic contexts. To the best of our knowledge, its association with an *Alphaproteobacteria* host has not been reported previously.

As expected, the majority of the class 1 integron gene cassettes we detected are AMR related. The dihydrofolate reductase encoded by *dfrA5* confers high levels of trimethoprim resistance in Gram-negative bacteria ^63^. We found an association between the *dfrA5* gene cassette and a *Comamonadaceae*-family host (Figure 2C). The *blaOXA*_10_ gene encodes a class D beta-lactamase that hydrolyses the semi-synthetic antibiotic oxacillin ^64, 65^. In our dataset, the *blaOXA_10_* gene cassette was detected in a *Moraxellaceae*-family host (Figure 2D). Genes *qacF* and *qacG2* encode small multidrug resistance (SMR) efflux pumps that confer resistance to quaternary ammonium compounds (QACs) present in a wide range of disinfectants and detergents ^31, 66^. We found a *qacG2* gene cassette in an *Aquabacterium*-like host (Figure 2E) and a *qacF* gene cassette at the proximal position in the gene cassette array in a *Rhizobacter* host (Figure 2F). Also found in the class 1 integron of the *Rhizobacter* host is an *aadA2* gene cassette. *aadA* genes confer resistance to the aminoglycoside antibiotics streptomycin and spectinomycin ^34, 67–70^. In the *Rhodocyclaceae*-family host, we found the same *aadA2* gene cassette with a SNP in the non-coding region and a non-synonymous mutation (V257A) in the coding region (Figure 2G and Table S3). The phylogenetic relationship between the seven hosts and the distribution of class 1 integron gene cassettes amongst them are summarised in Figure 3.

**Figure 3.**
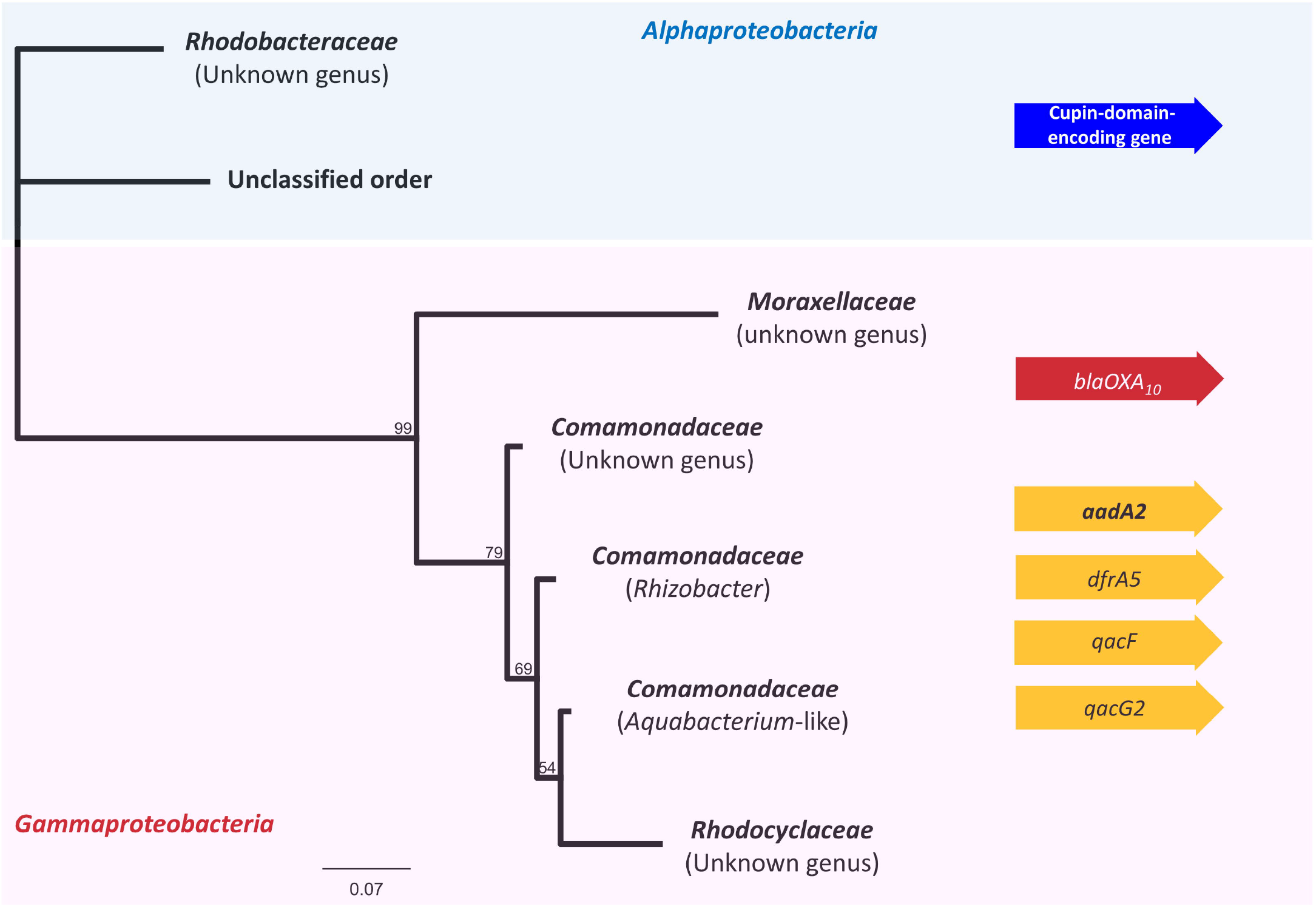
Phylogenetic tree constructed using the maximum likelihood method for the V4 hypervariable regions of 16S rRNA genes for the seven hosts of class 1 integrons identified in this study. The integron gene cassettes associated with *Alphaproteobacteria* (in blue) and *Gammaproteobacteria* (in red) are grouped accordingly. Gene cassettes that were found in *Gammaproteobacteria* hosts of the *Burkholderiales* order are shown in yellow.

### Discovery of *Rhizobacter* as novel carriers of class 1 integrons

*Rhizobacter* is an environmental bacterial genus. Isolates of *Rhizobacter* species have been successfully isolated from the rhizospheres of the *Panax* ^71, 72^ and *Bergenia* genera ^73^, as well as from freshwater sediment ^74^. There are still many questions regarding their potential ecological roles. For example, *R. gummiphilus* naturally degrades rubber ^44, 75, 76^, which implies beneficial roles in promoting nutrient recycling and bioremediation in soil. On the other hand, *R. daucus* is thought to be an agricultural plant pathogen that causes galls in carrot root tissues ^77^. Associations between the *Rhizobacter* genus and class 1 integrons have not been reported previously. In all currently available NCBI RefSeq genomes for the *Rhizobacter* genus (*n* = 11), we detected various non-class 1 integrons but did not find any bioinformatic evidence of class 1 integrons.

We found two *Rhizobacter* isolates with an identical nucleotide sequence in the 16S rRNA V4 hypervariable region to that of the *Rhizobacter* host in our dataset (NCBI accession no. AB835065.1 and LC734193.1) ^78, 79^. To experimentally verify that the association between the *Rhizobacter* and class 1 integron was *bona fide*, we designed *Rhizobacter*-specific 16S rRNA gene primers that target the same primer binding sites in the 16S rRNA gene as R926 and R806. By replacing the *intI1*_nested primer with *qacE*_F (the primer positions are indicated in Figure S3A) in our epicPCR experimental pipeline, we successfully obtained the expected chimeric epicPCR product featuring a *qacE* fragment and the *Rhizobacter* phylogenetic marker. This provides the first experimental evidence of an association between class 1 integrons and the *Rhizobacter* genus.

A recent study applied a modelling approach to understand the emergence and spread of novel antimicrobial resistance genes ^80^. The dispersal of antibiotic resistance genes from environmental bacteria to humans has been highlighted as a major knowledge gap that contributes to uncertainties in these models. Acquiring more experimental data for these gene mobilisation and HGT events can improve future quantitative risk assessment models for antibiotic resistance and our ability to curb transfer of antibiotic resistance genes between bacteria. epicPCR has the potential to contribute to this endeavour given its strengths in identifying environmental microbes that cannot be cultivated under laboratory conditions.

The seven bacterial hosts described in this study are expected to be a subset of all the class 1 integron hosts in our samples. Our bridging primer R519-*qacE* cannot bind to class 1 integrons that lack *qacE*/Δ*qacE*. These include many chromosomal class 1 integrons and atypical class 1 integrons with IS*26*-mediated deletions in their 3’-CS ^29, 41^. To detect these class 1 integrons and their hosts requires new bridging primers that bind to the 3’-end of their gene cassette arrays. In addition, the presence of epicPCR products that were detected in fewer than 3 replicates in this study could be confirmed by replacing the two *intI1*-targeting forward primers (*intI1*_outer and *intI1*_nested) with specific primers that target the 5’-end of specific gene cassettes of interest. The degenerate bases in the 16S rRNA gene-targeting primers also mean that certain hosts are less likely to be detected than others due to primer bias. Another inherent limitation is the variable resolution of taxonomic classifications when using 16S rRNA as phylogenetic marker.

We established epicPCR as a valuable surveillance platform for uncovering novel associations between class 1 integrons with diverse cassette arrays and their bacterial hosts in environmental water samples. Using this single-cell genomic approach, we successfully identified *Alphaproteobacteria* as novel hosts of a gene cassette encoding a cupin-domain protein of unknown functions, and the *Rhizobacter* genus as novel carriers of class 1 integrons. It is envisioned that this technology can be deployed to analyse water samples from sewage treatment plants, as well as effluents from industrial, clinical, and agricultural settings where antimicrobial usage is high. These surveillance efforts will hopefully lead to a timelier detection of hotspots of class 1 integron mediated AMR transmission and more targeted environmental mitigation measures. epicPCR also paves the way for future studies that seek to uncover the “sources” and “sinks” of integron gene cassettes in bacterial communities, as well as possible barriers to integron-mediated HGT that may also exist ^81^. With greater diversity in combinations of gene cassettes and bacterial hosts that are likely to be captured through future epicPCR studies, we will be able to address outstanding questions and gain more insights into possible origins of novel gene cassettes, which have so far remained the greatest mystery in the field of integron research ^32, 35, 82^.

## Associated Content

### Supporting Information

Additional experimental results and methods on the assignment of taxonomy for bacterial hosts (Table S1); oligonucleotide primers (Table S2); nucleotide sequences of gene cassettes (Table S3); hosts of the cupin-domain-encoding gene cassette (Table S4); sampling location and photograph (Figure S1); examples of fluorescence microscopy images for cell density estimation (Figure S2); additional epicPCR amplicons (Figure S3); nucleotide sequence alignments of conserved regions in sequenced class 1 integrons (Figure S4) (PDF)

### Author Contributions

Conceptualisation of study: TMG, QQ, SGT, MRG

Experimental investigation and validation: QQ, AP, JACS

Bioinformatic analysis: TMG, QQ, VR

Project management and supervision: MRG, SGT

Writing the original draft of the manuscript: QQ

Reviewing and editing the manuscript: All authors

### Funding Sources

This work was funded by the Australian Research Council Discovery Project DP200101874 grant to MRG and SGT.

## Acknowledgement

The authors thank Professor Marko Virta and his research group at the Department of Microbiology, University of Helsinki for sharing their expertise on epicPCR. We would like to acknowledge the Macquarie University Faculty of Science and Engineering Microscope Facility for access to its instrumentation and staff. We also thank Mr Richard Miller (IT Systems and HPC, Macquarie University) for his technical advice and support.

