## Supporting Information for "Uncovering bacterial hosts of class 1 integrons in an urban coastal aquatic environment with a single-cell fusion-polymerase chain reaction technology"

**Supplementary Tables**: Tables S1, S2, S3 and S4.

**Supplementary Figures**: Figures S1, S2, S3 and S4.

**Supplementary Files**: Available from the Dryad Data Repository https://doi.org/10.5061/dryad.b5mkkwhh4

**Table S1**: Assignment of taxonomy for bacterial hosts of class 1 integrons identified in this study using the SILVA ribosomal RNA gene database. The numbers in parentheses show nucleotide identity in the lowest classifiable taxonomic rank. Abbreviations: **c:** class, **o:** order, **f:** family, **g:** genus.

| Bacterial host | SILVA taxonomic classification |
|---|---|
| *Alphaproteobacteria* (Unclassified order) | **c:** *Alphaproteobacteria* (0.881) |
| *Rhodobacteraceae* (Unknown genus) | **c:** *Alphaproteobacteria*<br>**o:** *Rhodobacterales*<br>**f:** *Rhodobacteraceae* (0.988) |
| *Comamonadaceae* (Unknown genus) | **c:** *Gammaproteobacteria*<br>**o:** *Burkholderiales*<br>**f:** *Comamonadaceae* (0.988) |
| *Comamonadaceae* (*Aquabacterium*-like) | **c:** *Gammaproteobacteria*<br>**o:** *Burkholderiales*<br>**f:** *Comamonadaceae* (1.00) |
| *Comamonadaceae* (*Rhizobacter*) | **c:** *Gammaproteobacteria*<br>**o:** *Burkholderiales*<br>**f:** *Comamonadaceae*<br>**g:** *Rhizobacter* (1.00) |
| *Rhodocyclales* (Unknown genus) | **c:** *Gammaproteobacteria*<br>**o:** *Burkholderiales*<br>**f:** *Rhodocyclaceae* (0.996) |
| *Moraxellaceae* (Unknown genus) | **c:** *Gammaproteobacteria*<br>**o:** *Pseudomonadales*<br>**f:** *Moraxellaceae* (0.956) |

**Table S2:** Oligonucleotide primers used in this study.

(3SpC3 = 3' C3 spacer)

| Primer name | Oligonucleotide sequence (5’→3’) | Source |
|---|---|---|
| R926 | CCGYCAATTYMTTTRAGTTT | Parada et al (2016) |
| *intI1*_outer | CTGTAATGCAAGTAGCGTATGCG | This study |
| *intI1*_nested | CGAACGCAGCGGTGGTAA | |
| R519-*qacE* bridging primer | GWATTACCGCGGCKGCTGGCGGAAGGGGCAAGCTTAGT | |
| AP27_short | GCTCTTCCGATCTGGACTACHVGGGTWTCTAAT | Adapted from Diebold et al (2021) |
| AP28_short | GCTCTTCCGATCTGGACTAC | |
| Forward blocking primer | TTTTTTTTTTGWATTACCGCGGCKGCTG/3SpC3/ | Spencer et al (2016) |
| Reverse blocking primer | TTTTTTTTTTCAGCMGCCGCGGTAATWC/3SpC3/ | |
| *qacE*_F | GCAATAGTTGGCGAAGTAAT | This study |
| *Rhizobacter*_16S_ R926 | CCGTCAATTCCTTTGAGTTT | |
| *Rhizobacter*_16S_R806 | GGACTACCAGGGTATCTAAT | |

**Table S3:** Nucleotide sequences of all six unique gene cassettes that were detected by epicPCR. The coding region is underlined, while nucleotides of the *attC* sites are in bold. SNPs relative to the specified NCBI reference gene cassette sequences are shown in red.

| **Gene cassette** | **Nucleotide sequence (5' -> 3')** | **Comparison to NCBI reference gene cassette** |
|---|---|---|
| Cupin-domain-encoding gene | **TTAGAC**TTTTGGGCGGCGCGAATCTTGCTGCCCAAAGGCTTGCATTTACG<br>GAGCAAAAAATGAGCACTCACCGCGTGTATAAGTCCCATGAATTCATGC<br>AGCCAGCCGAGAAAGAACCAATACGGTCTGTTGTTGTTGAATCTTCGCAT<br>TCAGTTATCGTGGCTTGGCATGTGGAGCCCGGCCAAACTATTGCGCCACA<br>TATTCACCCCGACGGACAAGATACATGGACCATCCTTTCAGGGGAAGGT<br>CAATATCAAATTGACGAGCAAGGCAACACCGTTGCAATTGTTCCGGGGG<br>ATATCGTAGTTGCAAACAAAGGGCAGGTGCACGGTGTTCTTTGCACCAG<br>CGTTGGCCCACTTCGGTTCATTTCAGTGGTTGCGCCTTTGGATGCTGGTTA<br>TGAACCGCTATCAGCAAAAACATAGGGATATCAAGTGCAACAAGCTGC<br>CTCCGCTCCCCA**GCCTAACTGTTTGGTCAAGCTGACGCCAACAGGCTC**<br>**CGCCTGTTGGTACCCTCCGCTTCGCTCCGGCGCAGCTTACCGCGGG**<br>**CG** | 1 synonymous mutation in the coding region relative to the cupin-domain-encoding gene cassette in FJ172388 |
| *dfrA5* | **TTAAC**CCGGAACCAAAATTGTGAAAGTATCATTAATGGCTGCAAAAGCG<br>AAAAACGGAGTGATTGGTTGCGGTCCACACATACCCTGGTCCGCGAAAG<br>GAGAGCAGCTACTCTTTAAAGCCTTGACGTACAACCAGTGGCTTTTGGTG<br>GGCCGCAAGACGTTCGAATCTATGGGAGCACTCCCTAATAGGAAATACG<br>CGGTCGTTACTCGCTCAGCCTGGACGGCCGATAATGACAACGTAATAGT<br>ATTCCCGTCGATCGAAGAGGCCATGTACGGGCTGGCTGAACTCACCGAT<br>CACGTTATAGTGTCTGGTGGCGGGGAGATTTACAGAGAAACATTGCCCA<br>TGGCCTCTACGCTCCATATATCGACGATTGATATTGAGCCGGAAGGAGAT<br>GTTTTCTTTCCGAATATTCCCAATACCTTCGAAGTTGTTTTTGAGCAACAC<br>TTTAGCTCAAACATTAACTATTGCTATCAAATTTGGCAAAAG**GGTTAAC**<br>**AAAGCTATGCAATTGACGGTAAAAAGCTTCGTTCGCTTCGCTTGCTA**<br>**CGCTTCTTACCGCAATTGATAACGGCG** | 100% nucleotide identity with *dfrA5* gene cassette in X12868 |
| *blaOXA*$_{10}$ | **TTAGCC**ACCAAGAAGGTGCCATGAAAACATTTGCCGCATATGTAATTAT<br>CGCGTGTCTTTCGAGTACGGCATTAGCTGGTTCAATTACAGAAAATACGT<br>CTTGGAACAAAGAGTTCTCTGCCGAAGCCGTCAATGGTGTCTTCGTGCTT<br>TGTAAAAGTAGCAGTAAATCCTGCGCTACCAATGACTTAGCTCGTGCATC<br>AAAGGAATATCTTCCAGCATCAACATTTAAGATCCCCAACGCAATTATCG<br>GCCTAGAAACTGGTGTCATAAAGAATGAGCATCAGGTTTTCAAATGGGA<br>CGGAAAGCCAAGAGCCATGAAGCAATGGGAAAGAGACTTGACCTTAAG<br>AGGGGCAATACAAGTTTCAGCTGTTCCCGTATTTCAACAAATCGCCAGA<br>GAAGTTGGCGAAGTAAGAATGCAGAAATACCTTAAAAAATTTTCCTATG<br>GCAACCAGAATATCAGTGGTGGCATTGACAAATTCTGGTTGGAAGGCCA<br>GCTTAGAATTTCCGCAGTTAATCAAGTGGAGTTTCTAGAGTCTCTATATT<br>TAAATAAATTGTCAGCATCTAAAGAAAACCAGCTAATAGTAAAAGAGGC<br>TTTGGTAACGGAGGCGGCACCTGAATATCTAGTGCATTCAAAAACTGGTT<br>TTTCTGGTGTGGGAACTGAGTCAAATCCTGGTGTCGCATGGTGGGTTGGG<br>TGGGTTGAGAAGGAGACAGAGGTTTACTTTTTCGCCTTTAACATGGATAT<br>AGACAACGAAAGTAAGTTGCCGCTAAGAAAATCCATTCCCACCAAAATC<br>ATGGAAAGTGAGGGCATCATTGGT**GGCTAACAAGTCGCTCAAGGTCGC**<br>**TCCCTGCGGTCGCTGGACAGTCCCAGTCGGCGCATGCTTCGCATTTT**<br>**ATGCGCCGCCTGTGCCTGCCCCTTAGCTCCAACG** | 100% nucleotide identity with *blaOXA*$_{10}$ gene cassette in XXU37105 |
| *qacG2* | **TTAGAT**GCTTTGCTGTGCGCACAAATTTCGGCCAGCAACAAGACTGTTTT<br>TTTTCTTAAATCGAACCTAAAATTTCTTCGCGGAACTCCATGGAGAAATA<br>TTTTGAAAAATTGGTTATTTCTGGCTACGGCCATTATTTTTGAGGTCATTG<br>CAACCTCTGCGCTCAAGTCTAGTGAGGGCTTTACTAGGTTAGTACCGTCT<br>TTTATCGTCGTAGCGGGATACGCTGCTGCTTTTTATTTCCTGTCGCTGACA<br>CTCAAATCGATTCCTGTTGGAATCGCCTACGCAGTTTGGTCGGGCCTCGG<br>GATCGTCTTGGTCACTGCGATTGCATGGGTTTTGCATGGTCAAAAACTAG<br>ATATGTGGGGATTTGTTGGTGTCGGCTTCATTATCAGCGGCGTTGCTGTG<br>CTCAACTTGCTATCTAAGGCAAGTGTTCACTAAAACGGTCGC**ATCTAAC**<br>**CATTCCGTCGAGAGGGACCGCCCACAAGCTGCGCTTGCGGGTTCCC**<br>**TTCGCGGCTTCGCCGCTACGGCGGCCCCTCACGTCAAACG** | 2 non-synonymous mutations (S13F and A57G) in the coding region relative to *qacG2* gene cassette in AF327731 |

**Table S3 (continued)**

| Gene cassette | Nucleotide sequence (5' -> 3') | Comparison to NCBI reference gene cassette |
|---|---|---|
| *qacF* | **TTAGAT**GCCAGATTTGGCGTTGCGTATGCTCACAGAAAATCGATAGCC<br>GCAAGACTGTTTTTGGCAACTCATAGCCACTACAATTTCTCCTTCATAC<br>CGTAGAGGAGATTGCGCGTGAAGAACTGGATATTTCTGGCTGTTGCAA<br>TCTTTGGCGAGGTCATCGCAACTTCCGCACTGAAGTCTAGCCATGGATT<br>CACTAGGTTAGTTCCTTCCGTTGTAGTTGTGGCTGGCTACGGGCTTGCG<br>TTCTATTTCTTGTCTCTCGCGCTCAAGTCCATTCCGGTCGGTATTGCTTA<br>CGCTGTATGGGCTGGGCTTGGCATCGTGCTTGTGGCAGCTATTGCTTGG<br>ATTTTCCATGGCCAAAAACTAGACTTCTGGGCGTTCATTGGCATGGGAC<br>TTATCGTCAGTGGCGTCGCCGTTCTAAACCTGCTATCCAAGGTCAGCGC<br>ACATTGACCGGGTTGGC**ATCTAACAATTCATTCAAGCCGACGCCGCT<br>TCGCGGCGCGGCTTAATTCAGGCG** | 1 non-synonymous mutation (S10A) in the coding region relative to AF034958 |
| *aadA2*<br>(in *Rhizobacter* host) | **TTAGAC**ATCATGAGGGTAGCGGTGACCATCGAAATTTCGAACCAACTA<br>TCAGAGGTGCTAAGCGTCATTGAGCGCCATCTGGAATCAACGTTGCTG<br>GCCGTGCATTTGTACGGCTCCGCAGTGGATGGCGGCCTGAAGCCATAC<br>AGCGATATTGATTTGTTGGTTACTGTGGCCGTAAAGCTTGATGAAACGA<br>CGCGGCGAGCATTGCTCAATGACCTTATGGAGGCTTCGGCTTTCCCTGG<br>CGAGAGCGAGACGCTCCGCGCTATAGAAGTCACCCTTGTCGTGCATGA<br>CGACATCATCCCGTGGCGTTATCCGGCTAAGCGCGAGCTGCAATTTGG<br>AGAATGGCAGCGCAATGACATTCTTGCGGGTATCTTCGAGCCAGCCAT<br>GATCGACATTGATCTAGCTATCCTGCTTACAAAAGCAAGAGAACATAG<br>CGTTGCCTTGGTAGGTCCGGCAGCGGAGGAATTCTTTGACCCGGTTCCT<br>GAACAGGATCTATTCGAGGCGCTGAGGGAAACCTTGAAGCTATGGAAC<br>TCGCAGCCCGACTGGGCCGGCGATGAGCGAAATGTAGTGCTTACGTTG<br>TCCCGCATTTGGTACAGCGCAATAACCGGCAAAATCGCGCCGAAGGAT<br>GTCGCTGCCGACTGGGCAATAAAACGCCTACCTGCCCAGTATCAGCCC<br>GTCTTACTTGAAGCTAAGCAAGCTTATCTGGGACAAAAAGAAGATCAC<br>TTGGCCTCACGCGCAGATCACTTGGAAGAATTTATTCGCTTTGTGAAAG<br>GCGAGATCATCAAGTCAGTTGGTAAATGAT**GTCTAACAATTCGTTCA<br>AGCCGACCGCGCTACGCGCGGCGGCTTAACTCCGGCG** | 100% nucleotide identity with *aadA2* gene cassette in X68227 |
| *aadA2*<br>(in *Rhodocyclaceae* host) | **TTAGAC**ATCATGAGGGAAGCGGTGACCATCGAAATTTCGAACCAACT<br>ATCAGAGGTGCTAAGCGTCATTGAGCGCCATCTGGAATCAACGTTGCT<br>GGCCGTGCATTTGTACGGCTCCGCAGTGGATGGCGGCCTGAAGCCATA<br>CAGCGATATTGATTTGTTGGTTACTGTGGCCGTAAAGCTTGATGAAACG<br>ACGCGGCGAGCATTGCTCAATGATCTTATGGAGGCTTCGGCTTTCCCT<br>GGCGAGAGCGAGACGCTCCGCGCTATAGAAGTCACCCTTGTCGTGCAT<br>GACGACATCATCCCGTGGCGTTATCCGGCTAAGCGCGAGCTGCAATTT<br>GGAGAATGGCAGCGCAATGACATTCTTGCGGGTATCTTCGAGCCAGCC<br>ATGATCGACATTGATCTAGCTATCCTGCTTACAAAAGCAAGAGAACAT<br>AGCGTTGCCTTGGTAGGTCCGGCAGCGGAGGAATTCTTTGACCCGGTTC<br>CTGAACAGGATCTATTCGAGGCGCTGAGGGAAACCTTGAAGCTATGGA<br>ACTCGCAGCCCGACTGGGCCGGCGATGAGCGAAATGTAGTGCTTACGT<br>TGTCCCGCATTTGGTACAGCGCAATAACCGGCAAAATCGCGCCGAAGG<br>ATGTCGCTGCCGACTGGGCAATAAAACGCCTACCTGCCCAGTATCAGC<br>CCGTCTTACTTGAAGCTAAGCAAGCTTATCTGGGACAAAAAGAAGATC<br>ACTTGGCCTCACGCGCAGATCACTTGGAAGAATTTATTCGCTTTGTGAA<br>AGGCGAGATCATCAAGTCAGCTGGTAAATGAT**GTCTAACAATTCGTT<br>CAAGCCGACCGCGCTACGCGCGGCGGCTTAACTCCGGCG** | 1 SNP in the non-coding region, 1 non-synonymous mutation (V257A) in the coding region relative to X68227 |

**Table S4:** *Gammaproteobacteria* chromosomes and plasmids that contain the cupin-domain-encoding gene cassette, which shares 100% nucleotide identity with the NCBI reference gene cassette FJ172388.

| NCBI accession no. | Bacterial strain | Integron-integrase | Complete integron or CALINs |
|---|---|---|---|
| CP058131 (chromosome) & CP058136 (plasmid RHBSTW-00138_6) | *Klebsiella quasipneumoniae* strain RHBSTW-00138 | None | Chromosome and plasmid carry identical CALINs with 2 *attC* sites |
| LN879548 (Plasmid II) | *Comamonas thiooxydans* isolate C19 | IntI1 | Complete integron with 2 *attC* sites; cupin-domain-encoding gene cassette found at the proximal position of the gene cassette array |
| CP065406 | *Diaphorobacter* species JS3051 | 332 aa IntI; 58.3% pairwise aa identity to IntI1 | 1 complete integron with 14 *attC* sites; 1 CALIN with 10 *attC* sites |
| CP002657 | *Alicycliphilus denitrificans* strain K601 | 332 aa IntI; 59.4% pairwise aa identity to IntI1 | 1 complete integron with 8 *attC* sites; cupin-domain-encoding gene cassette found at the proximal position of the gene cassette array |
| AP024172 | *Alicycliphilus denitrificans* strain I51 | | 1 complete integron with 10 *attC* sites; 1 CALIN with 7 *attC* sites |
| CP043328 | *Pseudomonas aeruginosa* strain CCUG 51971 | IntI1 | 2 complete integrons with 5 *attC* sites in total |

#### Supplementary Figures

**Figure S1**: **(A)** Sampling location for coastal water at Shark Point, NSW, Australia **(B)** Photograph of rockpools downstream from a stormwater outlet at the sample collection site.

**Figure S2**: Representative images of DAPI-stained cells (in green) from coastal water samples that were imaged against 0.2 mm x 0.2 mm Neubauer grids on a haemocytometer by fluorescence microscopy.

**Figure S3**: **(A)** Correct fusion between class 1 integron and the V4 hypervariable region of 16S rRNA fragments should produce ~390 bp PCR products when epicPCR products are further amplified by the *qacE*_F and AP28_short primers. **(B)** To experimentally confirm *Rhizobacter* as *bona fide* hosts of class 1 integrons, we designed *Rhizobacter*-specific primers that bind to the V4 hypervariable region of 16S rRNA. The nucleotide sequences of the epicPCR product generated by the nested primers *qacE*_F and *Rhizobacter*_16S_ R806 were checked by Sanger sequencing.

**Figure S4**: Nucleotide sequence alignments of **(A)** the 157 bp *intI1-attI1* region in the 5'-CS region of class 1 integrons and **(B)** the 190 bp final *attC-qacE* region in the 3'-CS region of class 1 integrons that were sequenced in this study. With the single exception of a SNP and a single bp deletion that were found in a non-coding region of the class 1 integron associated with the *Rhodobacteraceae*-family host (indicated by the blue arrow), all the aligned nucleotide sequences were identical.

#### Supplementary Files

The sequencing data generated in this study are available from the Dryad Data Repository (https://doi.org/10.5061/dryad.b5mkkwhh4). The nine FASTQ files that are available for download are as follows:

- S1_1_Replicate_1.fastq
- S1_1_Replicate_2.fastq
- S1_1_Replicate_3.fastq
- S1_2_Replicate_1.fastq
- S1_2_Replicate_2.fastq
- S1_2_Replicate_3.fastq
- S1_3_Replicate_1.fastq
- S1_3_Replicate_2.fastq
- S1_3_Replicate_3.fastq

### Figure S1

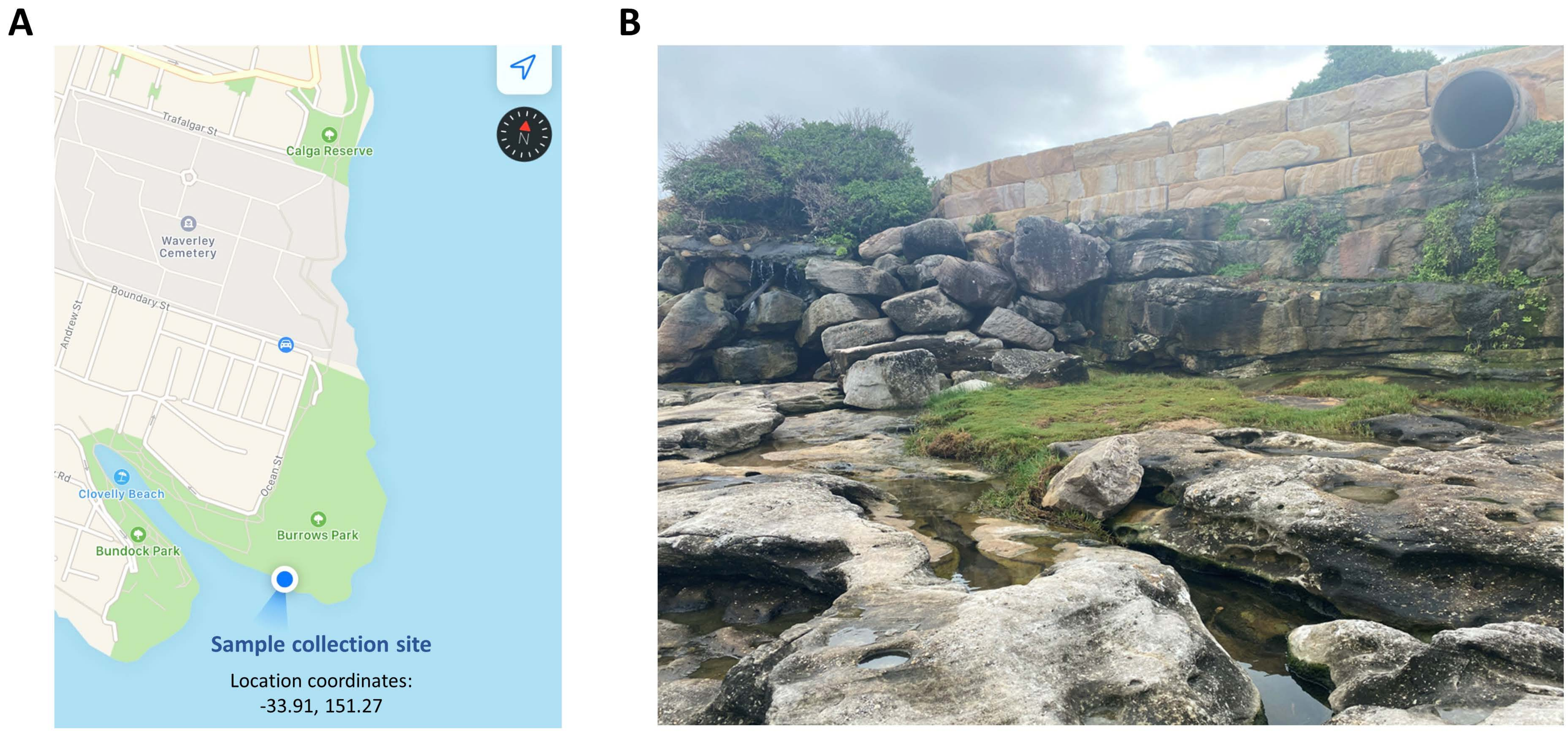
A
B
Trafalgar St
Calga Reserve
Waverley
Cemetery
Boundary St
Andrew St
Clovelly Beach
Ocean St
Burrows Park
Bundock Park
Sample collection site
Location coordinates:
-33.91, 151.27

### Figure S2

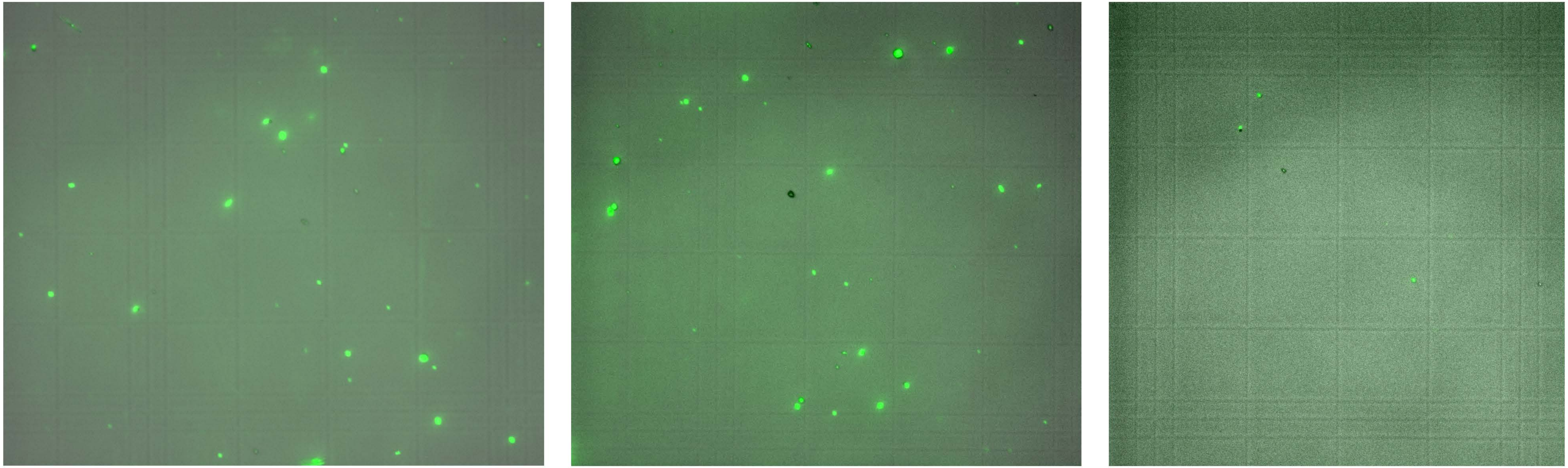

### Figure S3

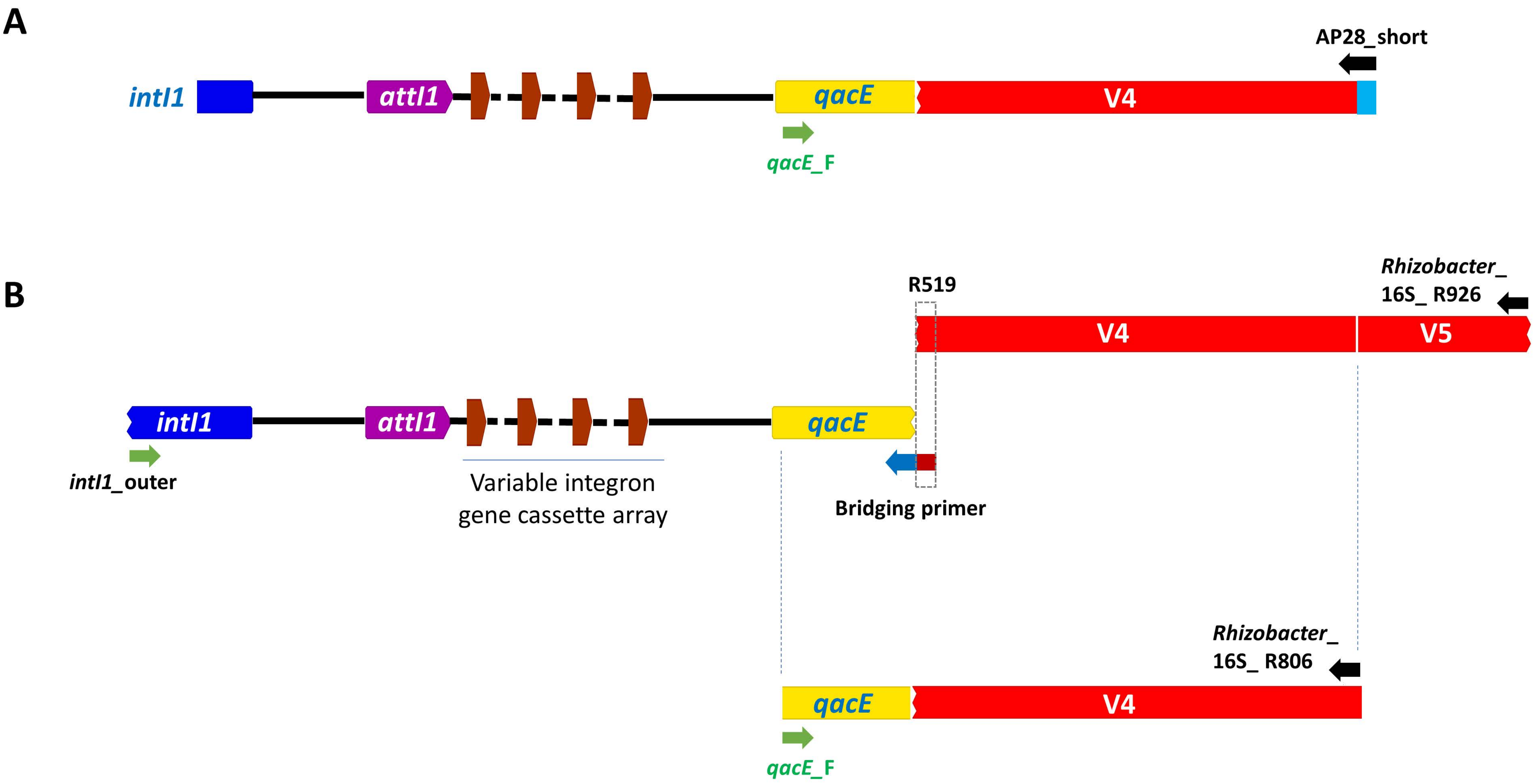
A
AP28_short
intI1
attI1
qacE
V4
qacE_F
B
R519
Rhizobacter_
16S_ R926
V4
V5
intI1
attI1
qacE
intI1_outer
Variable integron
gene cassette array
Bridging primer
Rhizobacter_
16S_ R806
qacE
V4
qacE_F

### Figure S4

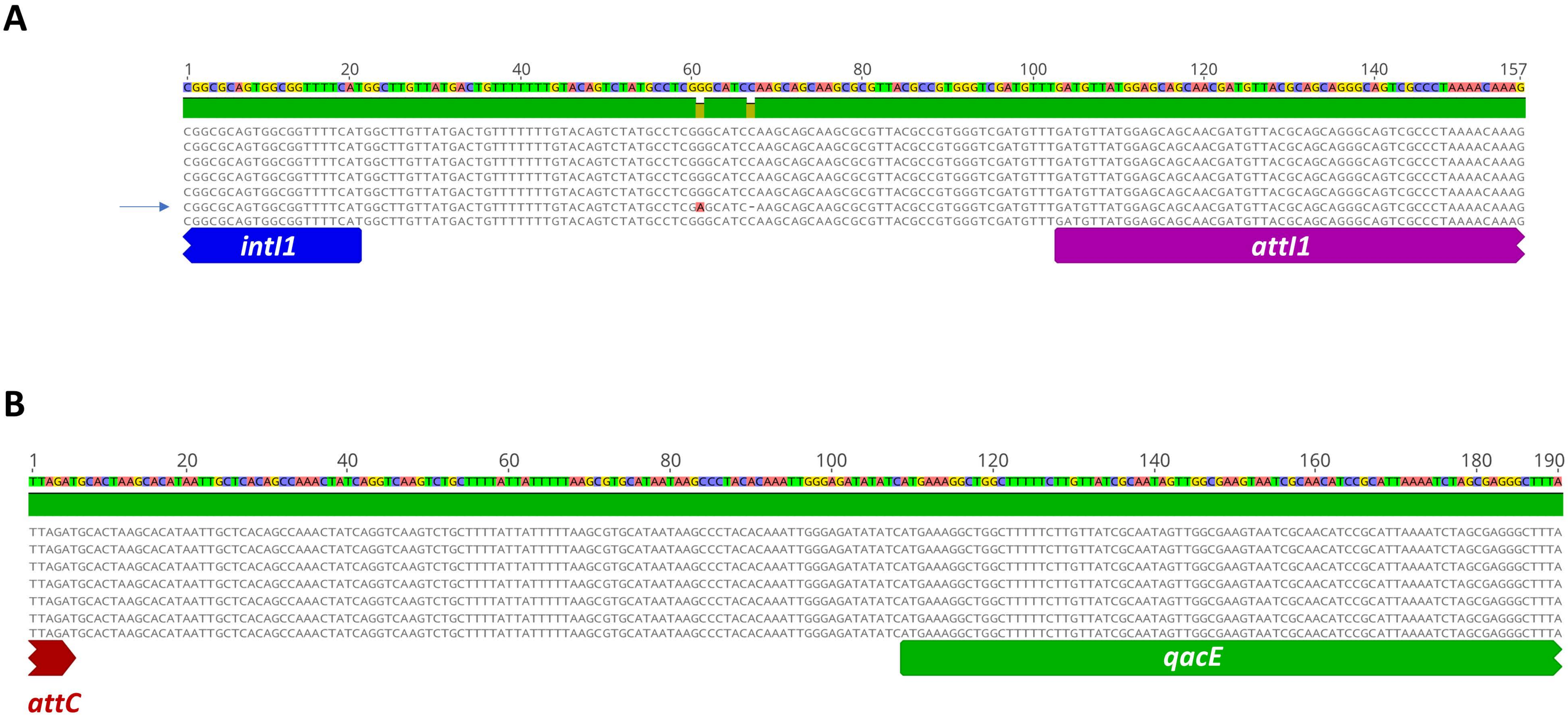
